# Skin protective effects of RM191A, a novel superoxide dismutase mimetic

**DOI:** 10.1101/2020.05.10.086694

**Authors:** Artur Shariev, Alistair J. Laos, Donna Lai, Sheng Hua, Anna Zinger, Christopher R. McRae, Llewellyn S. Casbolt, Valery Combes, Tzong-tyng Hung, Katie M. Dixon, Pall Thordarson, Rebecca S. Mason, Abhirup Das

## Abstract

Superoxide dismutase (SOD) is known to be protective against oxidative stress-mediated skin dysfunction. Here we explore the potential therapeutic activities of RM191A, a novel SOD mimetic, on skin. RM191A is a water soluble, dimeric copper (Cu^2+^-Cu^3+^)-centred polyglycine coordination complex. It displays 10-fold higher superoxide quenching activity compared to SOD as well as significant anti-inflammatory activity through beneficial modulation of several significant inflammatory pathways in cells.

We tested the therapeutic potential of RM191A in a topical gel using a human skin explant model and observed that it significantly inhibits UV-induced DNA damage in the epidermis and dermis, including cyclobutane pyrimidine dimers (CPD), 8-oxo-guanine (8-oxoG) and 8-nitroguanine (8NGO). RM191A topical gel is found to be safe and non-toxic in mice following month-long daily dosing at 0.19 mL/kg body weight. Moreover, it significantly accelerates excisional wound healing, and reduces 12-O-tetradecanoylphorbol-13-acetate (TPA)-induced skin inflammation in mice.

**Highlights:**

- Superoxide dismutase mimetic RM191A is a highly stable copper (Cu^2+^-Cu^3+^)-polyglycine coordination complex
- RM191A exhibits potent antioxidant (10-fold more than that of superoxide dismutase) properties *in vitro*
- RM191A exhibits potent anti-inflammatory properties *in vitro* and *in vivo*
- RM191A protects human skin explants against UV-induced oxidative stress and DNA damage
- RM191A is non-toxic and readily bioavailable in mice
- RM191A attenuates TPA-induced skin inflammation and improves wound healing in mice

## Introduction

Our skin acts as the first barrier of defence against various infections, trauma and environmental factors. Risk factors like ultraviolet (UV) radiation, environmental pollutants, xenobiotics, ageing, wounds, chronic inflammation and inflammatory skin diseases, such as psoriasis and atopic dermatitis, are major impediments against the proper maintenance of the skin barrier (1-4). Oxidative stress, which compromises proper skin integrity, is one of the most common features among these various risk factors (5-7). An over-abundance of oxidants or catalyst producing free radicals, such as reactive oxygen species (ROS) and reactive nitrogen species (RNS), and in parallel the failure of the endogenous antioxidant defence system in skin cells, primarily contribute to the formation of oxidative stress in skin (8-10). Furthermore, oxidative stress and inflammation are closely interconnected pathophysiological processes that are known participants in the pathogenesis of numerous skin diseases (11-14). Inhibition of oxidative stress is therefore important for maintenance of normal skin function.

Superoxide dismutase (SOD) enzymes catalyse the disproportionation of the cytotoxic superoxide (O_2_^-^) free radical to hydrogen peroxide and oxygen and play a significant role in the attenuation of oxidative stress in our body (15). SODs have been reported to be protective against many skin disorders (16). They sequester free radicals which are formed following UV exposure and have the potential to damage our DNA. Such DNA damage has been shown to be a major contributing factor to skin carcinogenesis. Due to its powerful antioxidant potential, endogenous SOD also exhibits potent anti-inflammatory properties by inhibiting the expression of ROS-sensitive transcription factors (17). SOD plays a major role in wound healing (18,19). For these reasons, exogenous SOD was contemplated as a therapeutic agent for the treatment of various skin disorders. However, the lack of clinical development of exogenous SOD is due to its large size (molecular weight ≈ 30 kDa), low cell permeability, short circulating half-life, antigenicity, and high manufacturing costs (20). To overcome these issues, much focus has been placed on the synthesis of low-molecular-weight, stable compounds that mimic SOD’s antioxidant activity.

There are three kinds of SOD enzymes. SOD2 is localized in mitochondria and contains manganese. Both cytosolic SOD1 and extracellular SOD3 enzymes contain a redox active copper atom (Cu^2+^), which is fundamental to the dismutation of the superoxide free radical, and a zinc atom (21-23). Interestingly, it has been established that if the zinc atom of either SOD is replaced with a copper atom, this di-copper form has the same free radical/superoxide scavenging activity as the endogenous enzymes (24). A significant number of the SOD mimetics reported in the literature focus on replicating the activity of SOD2. These include a variety of manganese porphyrin complexes, manganese (II) penta-azamacrocyclic complexes, and manganese (III) salen complexes (25). Topical application of these SOD mimetics demonstrated potential benefits for the treatment of skin disorders (26,27). Only a small number of SOD mimetics utilising the unique copper-based bivalent configuration of SOD1 have been reported, however, their biological activities were never demonstrated (24,28,29).

Here we report for the first time, a highly stable, unique copper (Cu^2+^-Cu^3+^)-centred polyglycine coordination complex – RM191A, which has been demonstrated through extensive analytical, *in vitro* and *in vivo* evaluation to possess robust antioxidant and anti-inflammatory activities. Our work indicates biological activity of topically applied RM191A across a broad spectrum of skin disorders. Specifically, we report protection of the epidermis, dermis and associated DNA following exposure to UV radiation in human skin explants; acceleration of wound healing; and significant anti-inflammatory and antioxidant activities in skin. We posit that these activities are a result of the complex’s SOD-like activity as well as its beneficial modulation of a number of important gene signalling pathways.

## Material and Methods

### RM191A Synthesis

RM191A was synthesized according to a proprietary protocol patented by RRMedSciences. RM191A gel formulation (5% RM191A hydrogel) consists of 21 mg of RM191A in 1 mL of hydrogel base.

### Elemental Analysis

Elemental microanalysis was performed using Perkin Elmer PE2400 elemental analyser (Macquarie University).

### FTIR

FTIR spectroscopy was performed using Nicolet IS5 FTIR Spectrometer at Macquarie University.

### Cryo-TEM

After diluting 2 mg of the sample into 1 mL of pure water (500 fold dilution), the sample (6 µL) was pipetted onto 300 mesh copper grids with a lacey formvar film (GSCu300FL-50, ProSciTech, Australia). The sample droplet was allowed to equilibrate for 30 seconds at room temperature and 90% relative humidity, before being blotted from one side for 1.8 seconds. The blotted grid was subsequently plunged into liquid ethane held at -174°C, excess ethane was blotted away with a piece of pre-cooled filter paper, and the vitrified grid stored in liquid nitrogen. The samples were then analysed by cryo-TEM at UNSW Mark Wainwright Analytical Centre.

### DOSY

The DOSY experiments were carried out on a Bruker Avance 400MHz NMR in D^2^O at 25°C with a BBFO probe using 3-9-19 water suppression at Mark Wainwright Analytical Centre, UNSW.

### X-ray crystallography

Blue solid RM191A was crystallised from supernatant generated by addition of ethanol to reaction mix. Powder diffraction was performed on a Panalytical Empyrean Bragg-Brentano geometry instrument fitted with a cobalt X-ray source at UNSW Mark Wainwright Analytical Centre.

### Measurement of SOD activity

We employed a modified version of the method described by Beauchamp and Fridovich (30). The objective was to create a reaction mixture that successfully generates known quantities of the O_2_^-^ free radical but does so at a pH at which RM191A is stable (pH 6.5 - 7.8). Nitro Blue Tetrazolium (NBT) was selected as an indicator to measure and compare the superoxide scavenging activities of both bovine SOD and RM191A. This assay utilized photochemical events to generate O_2_^-^ in a sodium tetraborate buffered solution of acetone and isopropanol which was then irradiated by high energy UV light at 254 nm (Mercury Vapour Lamp) to create O_2_^-^ in the solution. SOD and SOD-like substances inhibited the formation of the blue formazan by neutralising the superoxide free radicals as they were forming. Blue formazan has a characteristic UV absorbance at 560 nm. UV absorbances of the reaction solutions were measured at 560 nm following irradiation with UV light to determine the in situ superoxide free radical scavenging activity of both bovine SOD and RM191A. The UV-vis spectrum of both SOD and RM191A did not exhibit any absorbance at 560 nm (Fig. S2A). The reaction solution consists of 0.5 mL of 200 µmol/L NBT in phosphate buffer (pH = 7.8) and 0.5 mL of a 200 µmol/L solution of the test compound in 5.0 mL of 5 mol/L isopropanol. 15 mg acetone (20 µL) was added to begin the reaction. The reaction mixture was placed in UV light at 254 nm (mercury vapour lamp) for 1 minute and then transferred to a spectrometer to measure absorbance at 560 nm. The assay was repeated by adding, 1.0, 1.25, 1.5, 1.75, 2.0 mL of a 200 µmol/L solution of the test compound bringing total volume to 5.0 mL with 5 mol/L isopropanol.

### Cells

N27 cells were cultured in RPMI 1640 medium supplemented with 10% (v/v) fetal calf serum (FCS), 2 mM L-glutamine and penicillin/streptomycin (P/S) solution at 37°C with 5% CO_2_. RAW 264.7 cells were cultured in DMEM medium supplemented with 10% FCS and 1% P/S at 37°C with 5% CO_2_. Human umbilical vein endothelial cells (HUVEC) and hCMEC/D3 were cultured in RPMI + 10% FCS at 37°C with 3% O_2_ and 5% CO_2_. Adult normal human dermal fibroblasts (NHDF) were cultured in FGM-2 fibroblast growth medium-2 (Lonza) at 37°C with 3% O_2_ and 5% CO_2_.

### Cell viability

Cell viability was assessed using propidium iodide exclusion assay as described before (31). Briefly, cells (0.1 x 10^5^ cells) were seeded in a 48-well plate and incubated with different concentrations of RM191A or gel alone for 24 hrs. After 24 hrs, cells were trypsinized, stained with propidium iodide (PI, 10 µg/mL) and the number of PI negative (PI -ve) cells were determined by flow cytometry (BD FACSCanto II, Biological Resources Imaging Laboratory, UNSW). In hydrogen peroxide-mediated oxidative stress assay, N27 cells were first exposed to 200 µM hydrogen peroxide for 30 minutes and then treated with 21 µg/mL RM191A (gel as control) for 4 hrs, after which cell viability was measured using flow cytometry. Alternatively, cell viability was determined using MTT assay kit (Sigma) according to manufacturer’s protocol.

### LPS stimulation assay

LPS-stimulated ROS generation in RAW 264.7 cells was determined as previously reported (32). RAW 264.7 cells (10^6^) were stimulated with LPS (10 ng/mL) for 6 hrs with gel or RM191A (21 µg/mL) for 6 hrs. Fluorescent dye, CMH^2^DCFDA (final concentration 10 µL of 20 µM) was added to cells and incubated for 30 min in CO_2_ incubator to measure the intracellular ROS produced in these cells. After incubation, the cells were washed with cold phosphate buffered saline (PBS) and the amount of ROS was analysed by flow cytometry.

### Gene expression analysis

NHDF were treated with RM191A (0.5 mg/mL) or vehicle control. Mature RNA was isolated using an RNA extraction kit (Qiagen) according to the manufacturer’s instructions. RNA quality was determined using a spectrophotometer and was reverse transcribed using a cDNA conversion kit (Qiagen). The cDNA was used on the real-time RT^2^ Profiler PCR Arrays (QIAGEN, PAHS-065Y, PAHS-022Z and PAHS-162Z) in combination with RT^2^ SYBR® Green qPCR Mastermix (Qiagen). Fold changes in genes were calculated using delta delta C_T_ method in data analysis web portal (http://www.qiagen.com/geneglobe). Each sample was assayed in replicates and each group was analysed twice. The heat map was generated using Morpheus software (Broad Institute). The C_T_ cut-off was set to 36.

### Blood-brain-barrier transmission assay

hCMEC/D3 cell line derived from human brain endothelial cells were seeded onto membrane of 24 mm Transwells with 4 µm pores (Corning) coated with 3% collagen at 4.5 x 10^5^ cells/well and grown to confluence in RPMI + 10% FCS medium. RM191A was added to the top of the transwell at 0.02, 0.04, 0.08 and 0.4 mg/mL. After 2 and 6 hrs, media from top and bottom chambers of the transwells were collected and stored at -20°C until further analysis. The media was then analysed by HPLC (Phenomenex Luna HILIC Analytical Column 3 µm, 2 x 100 mm, mobile phase: 85/15 ACN/NH_4_ formate buffer (0.1M, pH 4.3)).

### UV exposure and RM191A treatment in skin explants

All experiments were performed according to procedures approved by University of Sydney Human Ethics Committee. Human skin explants were obtained with consent from patients undergoing elective surgery. The protocol for obtaining skin explants and for their use to examine photoprotective effects of test agents was described in Song *et al.*, with some minor changes (33). In brief, skin removed at elective surgery was cleaned, subcutaneous fat removed and cut into pieces to produce multiple pieces approx. 4-6 mm x 4-6 mm, with each explant placed in one well of a 96 well plate in RPMI 1640 media containing 10% FCS. Skin explants in triplicate were used for each treatment point. The DNA damage was analysed after 3 hrs incubation period.

Skin was subjected to solar simulated UV-radiation, as previously described (33,34). The spectral output of this solar simulator compared with sunlight is shown in Fig. S3A. Treatments, as indicated below, were applied to the epidermal surface of the explant. Sham wells were treated in a similar manner to UV-exposed wells, including media changes and treatments, but were covered with foil and not exposed to UV. RM191A gel or the control gel was used with two protocols. In Protocol 1, 1.5 µL of RM191A or gel was applied to the surface of the explant 10-15 minutes prior to UV exposure, then again immediately after UV and then at 0.5, 1.5, and 2.5 hrs. In Protocol 2, 1.5 µL of RM191A or gel was applied to the surface of the explant immediately after UV exposure, then at 1, 2, 2.5 hrs. In addition, vehicle containing wells consisted of RPMI media with 10% v/v FCS and 0.1% v/v spectroscopic ethanol applied immediately after UV. The positive control treatment was 1,25 dihydroxyvitamin D3 (1,25D) (33), which was dissolved in spectroscopic grade ethanol at a final concentration of 1 nM and 0.1% v/v ethanol in RPMI media with 10% v/v FCS. This was also applied immediately after UV only. Sham wells consisted of vehicle (containing ethanol 0.1% - immediately after UV exposure), 1,25D (applied immediately after UV exposure), Gel 1 (Protocol 1 only) and RM191A 1 (Protocol 1 only). There were insufficient explants to do sham wells of RM191A and gel with Protocol 2. UV-exposed wells consisted of vehicle, 1,25D, Gel 1, Gel 2 (Protocol 2 only), RM191A 1 and RM191A 2 (Protocol 2 only).

### Animal experiments

All experiments were performed according to procedures approved by UNSW Animal Care and Ethics Committee (UNSW, Australia). Male C57BL/6J mice (Australian BioResources, Mossvale, NSW and Animal Resource Centre, Canning Vale, WA, Australia) were housed in the UNSW Biological Resource Centre at 21°C ± 2, with a 12 hrs/12 hrs dark/light cycle and were fed standard chow diet.

### RM191A short-term and long-term treatment

One day prior to the application of RM191A or gel, male mice were anaesthetised and hair on the dorsal surface was removed with a clipper. The shaved area was then cleaned with betadine solution, wiped with an alcohol swap and dried for a few seconds. The following day, the animals were briefly anaesthetised and 50 µL of RM191A at different concentrations was applied topically. This step was repeated for 4 and 29 consecutive days for short-term and long-term treatment protocols, respectively. On the 4^th^ and 30^th^ day, the animals were euthanized, tissues were harvested, and flash frozen in liquid nitrogen. The overall health of the animals was monitored daily and weights were recorded daily until euthanized. For the short-term treatment protocol, the treatment groups consisted of gel and RM191A (0.19, 0.38, 0.76, 1.33 and 1.9 mL/kg body weight). For short-term treatment protocol, the groups consisted of no gel, gel and RM191A (0.19 mL/kg body weight).

### Rotarod test

Motor coordination was assessed using the rotarod test as previously described (35). In short, mice were acclimatized to a rotarod for 3 days in three trials lasting 2 min each at a constant speed of 5 rpm. On the 4th day, the animals were subjected to three trials on the accelerating roller (4 - 40 rpm in 4 min) and the time that the mice remained was recorded.

### Rearing test

Mouse rearing test was performed as described previously (36). Briefly, the mouse was placed in a clean transparent cylinder (diameter: 12 cm, height: 15 cm) for 3 minutes and its forelimb activity while rearing against the wall of the arena was recorded.

### EchoMRI

Body composition (fat mass, lean mass, and total body water) was measured by EchoMRI (EchoMRI-900, Mark Wainwright Analytical Centre, UNSW).

### RM191A time point study

One day prior to the application of RM191A in mice, the animals were anaesthetised and hair on the dorsal surface was removed with a clipper. The shaved area was then cleaned with betadine solution, wiped with alcohol swap and dried for a few seconds. Next day, the animals were briefly anaesthetised and 50 µL of RM191A in gel formulation (1.9 mL/kg body weight) was applied topically. The animals were euthanized at 15 min, 30 min, 1 hr and 2 hrs post-treatment and tissues were harvested. Untreated animals were used for t = 0.

### ICP-MS

About 100 mg of tissue sample (50 µL in the case of plasma and urine) was transferred to a 10 mL polypropylene tube (Sarstedt), then 200 µL of concentrated nitric acid (HNO_3_) was added and incubated for 12 hrs at room temp with occasional shaking. After 12 hrs, the tube was heated at 80°C moderately up to clearing (5 hrs) and cooled down to room temperature. Then 20 µL of hydrogen peroxide was added, and the tube was again heated upto 80°C for 2 hrs. Next, the volume was made up to 10 mL with milliQ water. A blank standard was prepared in the same way without tissue samples. The samples were then analyzed for copper using ICP-MS (PerkinElmer Nexicon) at Mark Wainwright Analytical Centre, UNSW.

### Ear edema model

TPA (12-O-tetradecanoylphorbol-13-acetate)-induced mouse ear oedema was generated according to a published protocol (37). Briefly, 6 µg of TPA dissolved in 20 µL acetone was applied to the left ear of a mouse. Acetone (20 µL) was applied to the right ear as vehicle control. Both ears were pre-treated with 50 µL of gel or RM191A (21 µg/mL), 15 minutes prior to TPA or acetone application. After 6 hrs, ear tissue was excised using a 5 mm biopsy punch and ear weight was measured. The ear samples were then fixed in 10% formalin.

### Wound regeneration model

Wound healing protocol in mice was performed as described previously, with minor modifications (38). One day prior to wounding, the animals were anaesthetised and hair on the dorsal surface was removed with a clipper. The shaved area was then cleaned with betadine solution, wiped with an alcohol swab and dried for a few seconds. Next day, the animals were briefly anaesthetised, and a wound was created on the dorsal skin using a 10 mm biopsy punch. The picture of wound was captured, and the wound was covered with a semi-occlusive dressing (3M Tegaderm). Two days after the wounding procedure, 50 µL of gel or RM191A (21 µg/mL) was applied to the wound area and the application was repeated every 2 days. Each animal was singly housed during the entire procedure. The pictures of wounds were captured every 2 days and wound area was analyzed using ImageJ software.

### Hematoxylin and eosin (H&E) staining

After fixation in paraffin, samples were stained with 0.1% Hematoxylin for 10 min, rinsed with deionised H_2_O, stained with Scott’s blue solution for 1 min and then washed with deionised H_2_O. The sections were then dipped in Eosin for 3 min, dehydrated through alcohol and cleared in xylene. The slides were mounted with DPX and imaged using a Zeiss Axioscan Z1 slide scanner (University of Sydney).

### Immunohistochemistry

Three or 24 hours after irradiation, the explants were fixed, embedded in paraffin and sectioned before being deparaffinized and rehydrated for staining. Antigen retrieval was achieved with 10 mM citrate buffer. These sections were then stained as described for cyclobutane pyrimidine dimers (CPD), oxidative DNA damage in the form of 8-oxo-guanine (8-oxoG) and nitrosative DNA damage in the form of 8-nitro-guanine (8NGO) (33). Isotype controls were performed (without primary antibody). Immunohistochemistry was carried out and analysed using the Metamorph image analysis program as previously described (33), except that the results were expressed as mean pixel intensity – mean gray value in arbitrary units (34). In addition to analysing nuclear staining in the epidermis, for 8-oxoG and 8NGO, a separate analysis was performed on the upper layer of the dermis, where there were reasonable numbers of cells visible.

### Statistics

Data are presented as mean ± SEM. Statistical significance was performed using Student’s t test or ANOVA with Tukey’s *post hoc*test. Statistical tests were performed using GraphPad Prism software. P values of less than 0.05 were considered statistically significant.

## Results

### Synthesis and characterization of RM191A

Synthesis of RM191A yielded a dark blue-green liquid which contains 11.8% copper (Fig. 1A, Table 1). RM191A was a highly polar complex that is completely insoluble in non-polar solvents. Although the UV-visible absorption spectra of RM191A was similar to that of Cu-EDTA complex (Fig. S1A), RM191A exhibited an intense fluorescence emission peak at 440 nm (Fig. 1B). FTIR spectra of RM191A showed absorbances characteristic of amino and carboxylic acid functionalities (Fig. S1B). This information combined with the elemental analysis indicated that RM191A is a copper-amino acid coordination complex.

**Figure 1:**
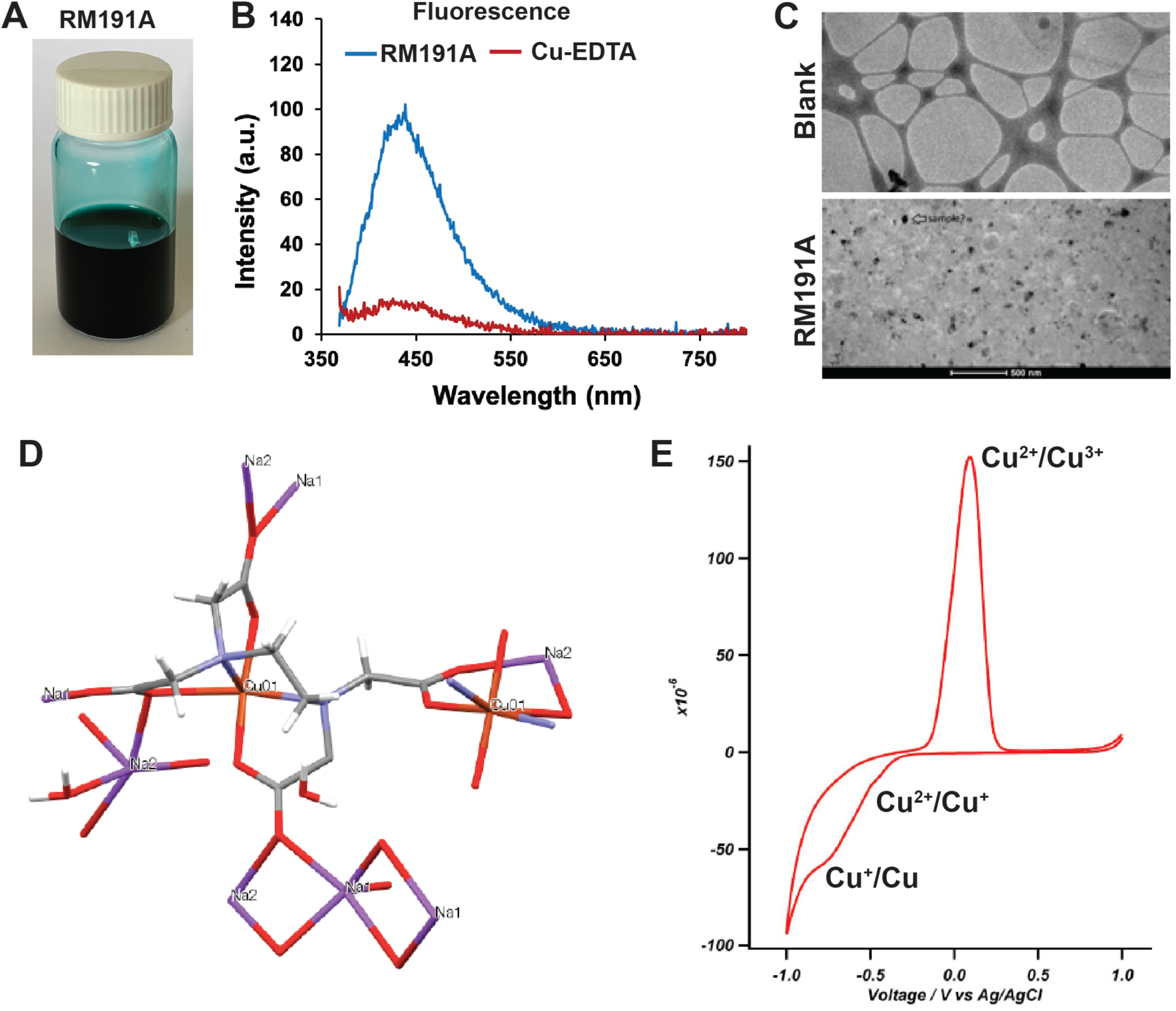
Synthesis and characterization of RM191A. (A) Picture of RM191A solution. (B) Fluorescence emission spectrum of RM191A and Cu-EDTA. (C) Cryo-TEM images of blank (top) and RM191A (bottom). (D) Structure of RM191A crystal from X-ray single-crystal analysis (Red = oxygen, blue = nitrogen, light red = copper, dark grey = carbon, light grey = hydrogen). (E) Cyclic Voltammetry results of RM191A showing two reduction peaks and one oxidation peak.

**Table 1:** Elemental analysis of RM191A.

| Element | Cu | C | H | N | Na | O |
| --- | --- | --- | --- | --- | --- | --- |
| % | 11.8 | 27.46 | 4.45 | 7.06 | 12.0 | 37.23 |

To delineate the structure further, analysis via cryo-TEM was performed in which RM191A showed unexpected black dots of about 20-50 nm in diameter in comparison with a blank sample, strongly suggesting the presence of large copper-containing complexes in this sample (Fig. 1C). This conclusion was also supported by NMN diffusion ordered spectroscopy (DOSY), in which the compound exhibited a large apparent increase in the chelate hydrodynamic radii suggesting that the proton signals seen in the ^1^H NMR of the copper chelate complex belong to a species that is oligomeric (Table 2, Fig. S1C). The hydrodynamic radius of Cu species in RM191A appeared to be at least 4.67 Å or a diameter of nearly 1 nm compared to 3.0 Å radius (0.6 nm diameter) for a standard Zn-EDTA complex. X-ray diffraction of crystalline RM191A showed a hydrated copper-polyglycine coordination complex consisting of two copper atoms with square planar and pyramidal geometry (Fig. 1D, S1D). Diffusion pulsed voltammetry confirmed the presence of both Cu^2+^ and Cu^3+^ ions in RM191A (Fig. 1E, S1E).

**Table 2:** DOSY analysis of Zinc-EDTA and RM191A complex.

| Compound | log(D) (m<sup>2</sup>S<sup>-1</sup>) | Hydrodynamic radius (m) | Hydrodynamic volume (m<sup>3</sup>) | Relative volumes |
| --- | --- | --- | --- | --- |
| Zinc-EDTA | -9.09 | 3.02 x10<sup>-10</sup> | 1.15 x10<sup>-28</sup> | 1 |
| RM191A | -9.28 | 4.67 x10<sup>-10</sup> | 4.27 x10<sup>-28</sup> | 3.7 |

### RM191A has potent antioxidant and anti-inflammatory properties

The interesting similarity of RM191A to the dimeric structure of SOD1 and presence of unique Cu^2+^-Cu^3+^ dipole led us to postulate that RM191A would be a highly efficient free radical scavenger which could act in a similar manner to SOD. To prove our hypothesis, we measured and compared the superoxide scavenging activities of both bovine SOD and RM191A using a well-established assay (30). The results showed that at 20 µM concentration, RM191A was 10 times more effective at neutralising superoxide free radicals than SOD (Fig. 2A). With increasing concentration, the activity of RM191A exceeded that of SOD by 30 times.

**Figure 2:**
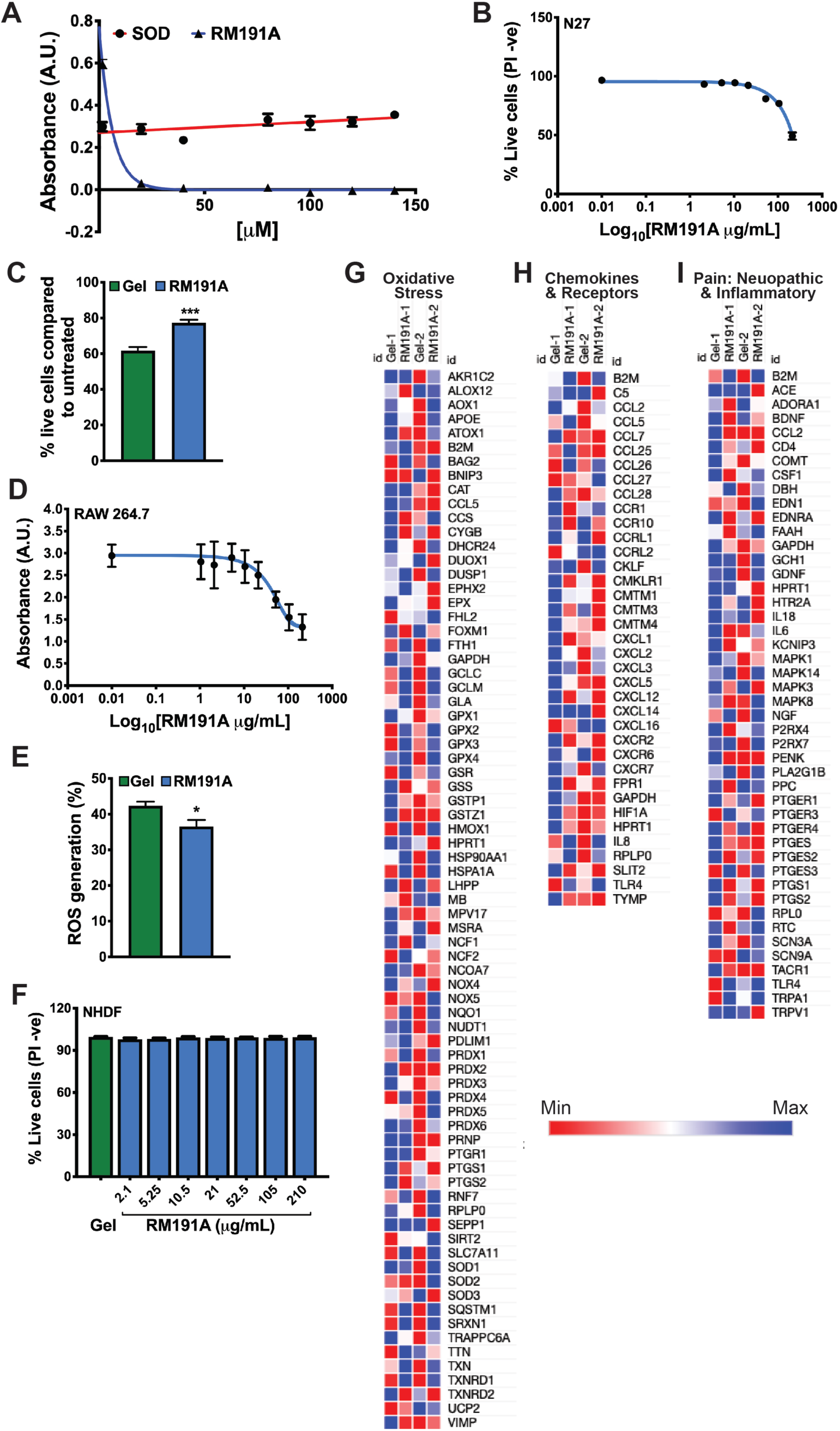
RM191A has potent antioxidant and anti-inflammatory properties. (A) Comparison of superoxide scavenging activities of both bovine SOD1 and RM191A (n = 3) at 560 nm. (B) Dose dependent curve of RM191A for N27 cells, as measured by PI exclusion (PI -ve) in flow cytometry (n = 6). (C) Percentage live N27 cells, as measured by PI exclusion (PI -ve) in flow cytometry, exposed to 200 µM hydrogen peroxide for 30 minutes and then treated with 21 µg/mL RM191A or gel for 4 hrs (n = 6). (D) Dose dependent curve of RM191A for RAW 264.7 cells, as measured by MTT assay (n = 8). (E) Amount of ROS generated by RAW 264.7 cells, as measured by CM-H_2_DCFDA fluorescence in flow cytometry, upon LPS (10 ng/mL) stimulation and followed by treatment with RM191A or gel only (n = 6). (F) Adult normal human dermal fibroblasts (NHDF) viability at different doses of RM191A, as measured by PI exclusion (PI -ve) in flow cytometry (n = 6). Heat map generated from the C_T_ values in PCR arrays reflecting (G) Oxidative Stress, (H) Chemokines and Receptors and (I) Pain-associated gene expression in NHDF treated with or without RM191A. The C_T_ cut-off was set to 36. Data expressed as mean ± SEM. *p < 0.05 and ***p < 0.0005 by t-test.

Next, we decided to test the bioactivity of RM191A in cells. For all subsequent experiments, RM191A was dissolved in a hydrogel (Gel). Upon treatment of N27 cells, an immortalized rat dopaminergic neuronal cell line (39), with different concentrations of RM191A and measuring the percentage of live cells by flow cytometry or MTT assay, the LD_50_ of RM191A was observed to be 210 µg/mL (Fig. 2B, S2B). Similarly, LD_50_ of RM191A for human umbilical vein endothelial cells (HUVEC) was found to be 210 µg/mL (Fig. S2C). In order to determine if RM191A could protect cells from ROS-mediated oxidative stress, N27 cells were first exposed to 200 µM hydrogen peroxide for 30 minutes and then treated with 21 µg/mL RM191A (gel as control) for 4 hrs. RM191A treatment increased the viability of cells by 24%, when measured by flow cytometry, compared to the control (Fig. 2C).

ROS are known to mediate inflammatory responses induced by a variety of stimuli including lipopolysaccharide (LPS) (40,41). Treatment of macrophage-like RAW 264.7 cells with SOD decreases LPS-induced ROS generation and upregulation of several inflammatory genes (41,42). To investigate if RM191A has similar effects, at first, we determined its toxicity in RAW 264.7 cells using flow cytometry and MTT assay. The LD_50_ of RM191A was 85 µg/mL in both assays (Fig. 2D, S2D). Next, RAW 264.7 cells were stimulated with LPS (10 ng/mL) and then treated with RM191A or gel only. The amount of ROS was measured by flow cytometry. LPS stimulation increased the ROS by 42%, however, treatment with RM191A decreased ROS levels to 36% (Fig. 2E). Given this data, we were curious to understand RM191A’s ability to affect oxidative stress and inflammatory pathways. To investigate this, adult normal human dermal fibroblasts (NHDF) were treated with RM191A and changes in gene expression compared to control were determined using PCR arrays. Toxicity analysis demonstrated that RM191A was well-tolearetd by NHDF, with no evident cell death upto a dose of 210 µg/mL (Fig. 2F). Although RM191A treatment did not change the expression of SODs in NHDF, the expression of several genes that confer cellular protection against oxidative stress were significantly upregulated (Fig. 2G, Table S1). Notably, heme oxygenase (HMOX1), heat shock protein (HSPA1A) and thioredoxin reductase (TXNRD1) expression increased by 67, 7 and 5-fold respectively in RM191A-treated NHDF compared to the control. Amongst the inflammatory genes, CCL7, ACKR4, CXCR6 and CMKLR1 were downregulated, while the expression of CCL26, CXCL8 and TLR4 were upregulated in RM191A-treated NHDF (Fig. 2H, Table S2). These small changes in chemokines may be due to the fact that in basal condition, the inflammatrory genes are not upregulated in NHDF and therefore the protective effects of RM191A were not obvious. Repeating the experiment where the NHDF are subjected to oxidative stress by external stimuli would be ideal to demonstrate the anti-inflamamtory effects of RM191A. Since inflammation also contributes to pain (43), we measured the effects of RM191A on pain-associated genes. Several key biomarkers of pain, for example, IL-18, CCL2, serotonin receptor (HTR2A) and ion channel (TRPV1) were downregulated in NHDF upon RM191A treatment (Fig. 2I, Table S3).

### RM191A protects skin against UV-induced oxidative stress and DNA damage

Exposure of skin to UV radiation induces direct DNA damage such as generation of cyclobutene pyrimidine dimers (CPD), as well as oxidative stress via a dramatic increase in ROS (44). Such an increase results in characteristic DNA damage, for example, oxidative DNA damage in the form of 8-oxo-guanine (8-oxoG) and nitrosative DNA damage in the form of 8-nitroguanine (8NGO) (45). If these DNA lesions are inadequately repaired, mutations may occur and may lead to the development of skin tumours (45). We wanted to determine whether RM191A, when applied topically to *ex vivo*human skin, reduced UV-induced DNA damage.

Human skin was either pre-treated with RM191A (RM191A 1), followed by exposure to UV radiation, or treated with RM191A (RM191A 2), immediately after UV exposure. In both cases, RM191A treatment post-UV exposure was repeated every 30 minutes for 3 hrs incubation period. The same treatment protocols were also followed for gel (Gel). An active form of vitamin D – 1,25-dihydroxyvitamin D3 (1,25D), was used as a positive control in these experiments as it has previously been shown that topical application of 1,25D reduces UV-induced DNA damage in human skin cells, human skin explants and human subjects, and reduces UV-induced skin tumours in mice (46,47). Ethanol (Vehicle) was used as a control for 1,25D treatment. Staining for CPD, was virtually absent in the isotype control, and the intensity of nuclear staining was low in the sham exposed explants, indicating very few CPDs (Fig. S3B). This was expected, since exposure to UV was the major source of the energy required to produce CPD. Exposure to UV significantly increased CPD in UV-vehicle or UV-gel groups compared with their respective sham controls (compare Fig. 3A and Fig. S3B). Treatment of UV-exposed explants with 1,25D or RM191A (RM191A 2) decreased CPD by 30% compared to their respective UV-exposed controls (Fig. 3A). When pre-treated with RM191A (RM191A 1), CPD decreased by 37% compared to its control (Fig. 3A).

**Figure 3:**
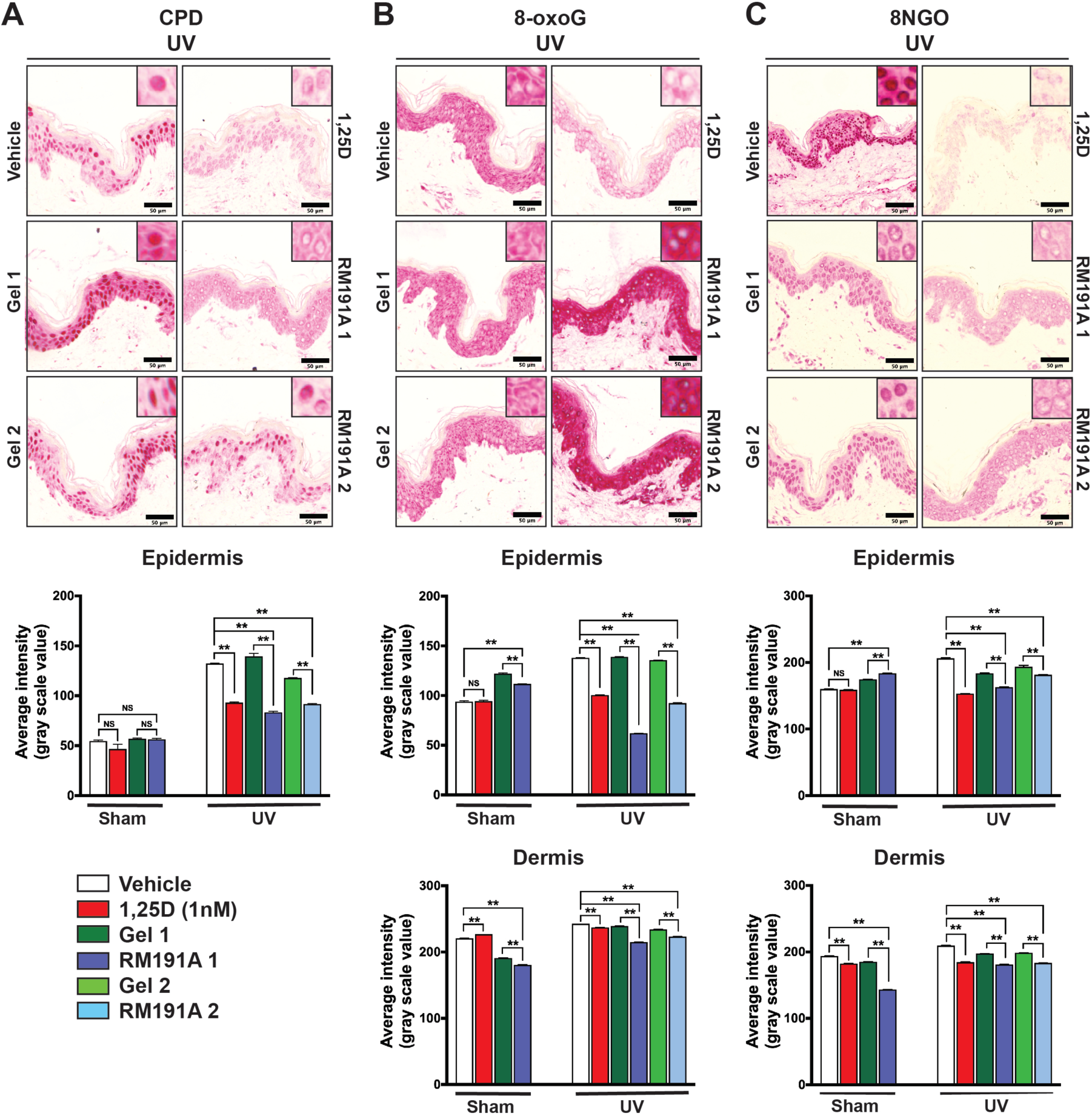
RM191A protects against UV-induced oxidative stress. (A) Representative images of immunohistochemical staining of CPDs in human skin explants from UV-exposed explants. Analysis of CPD in epidermis of skin explants. (B) Representative images of immunohistochemical staining of 8-oxoG in human skin explants from UV-exposed explants. Analysis of 8-oxoG in epidermis and dermis of skin explants. (C) Representative images of immunohistochemical staining of 8NGO in human skin explants from UV-exposed explants. Analysis of 8NGO in epidermis and dermis of skin explants. The maginified (120X) images of each stainings are shown in the insets. The red stain in the 8-oxoG is in the cytoplasm and the nuclei in the RM191A treated groups are considerably less stained. Data expressed as mean ± SEM. ** p < 0.001 and NS = not significant by One-way ANOVA between data sets (n = 3 explants per data point).

The intensity of nuclear 8-oxoG staining was moderate in the sham exposed explants treated with vehicle or 1,25D (Fig. S3C) due to the fact that culture of the explants engendered some oxidative stress. Markedly increased cytoplasmic staining was seen in explants which were sham exposed and treated with either gel or RM191A (Fig. S3C). Because of this high level of cytoplasmic staining, which was also observed in UV exposed explants, for the analysis, two separate masks (cytoplasmic and nuclear) were created using the Metamorph image analysis program and only the nuclear staining was imaged, so as not to include this strong background, non-specific cytoplasmic stain. Exposure to UV significantly increased nuclear 8-oxoG in both the UV-vehicle and UV-gel groups compared with their respective sham controls (compare Fig. S3C and Fig. 3B). Treatment of UV-exposed explants with 1,25D decreased nuclear 8-oxoG by 27% compared to their UV-exposed controls. Treatment with RM191A (RM191A 2) was found to decrease 8-oxoG levels by 33%, and the RM191A pre-treated group (RM191A 1) was found to have 55% less 8-oxoG, when compared to UV-exposed controls (Fig. 3B). Interestingly, oxidative DNA damage in the nuclei of epidermal cells was lower in RM191A-treated sham explants compared to gel-treated sham wells, indicating that RM191A was able to reduce even base-level oxidative damage. There were far fewer cells in the dermis of the explants, but an analysis was performed on the intensity of staining in the nuclei of cells of the upper dermis. Nuclear staining for 8-oxoG was overall lower in the dermis of sham exposed explants than in UV-exposed explants, but it was not negligible (Fig. 3B). Pre-treatment of sham explants with RM191A significantly reduced 8-oxoG staining. Nuclear stain for 8-oxoG was significantly increased in the dermis in vehicle and gel treated UV-exposed explants, compared with sham exposed controls, though the fold increase with UV was not as great as seen in the epidermis, probably reflecting reduced penetration of UV into the dermis. Nevertheless, as shown in Figure 3B, each of the active treatments significantly reduced 8-oxoG in the dermis.

Similar to 8-oxoG staining, exposure to UV significantly increased nuclear 8NGO staining in UV-vehicle and UV-gel groups, in both the epidermis and dermis, compared with their respective sham controls (compare Fig. S3D and Fig. 3C). As shown in Figure 3C, treatment of UV-exposed explants with 1,25D or with RM191A using either protocol significantly decreased 8NGO compared with their UV-exposed controls. Again, pre-treatment of UV-exposed explants with RM191A was more effective at reducing nitrosative DNA damage than RM191A post-treatment.

### RM191A is non-toxic and readily bioavailable in mice

After determining the biological activities of RM191A in cells and skin explants, we sought to determine if these effects can translate *in vivo.*Firstly, we used a mouse model to test if RM191A has any toxicity in animals. The dorsal surfaces of 12 week old male mice were shaved and 50 µL of RM191A at different doses (0.19 - 1.9 mL/kg body weight) was applied topically daily for 4 days. We found that there were no significant changes in the body weight or tissue weights of these animals compared to vehicle control (gel only) after 4 days of topical treatment (Fig. 4A, Table S4), implying that short-term treatment of RM191A is well tolerated in mice up to a concentration of 1.9 mL/kg body weight.

**Figure 4:**
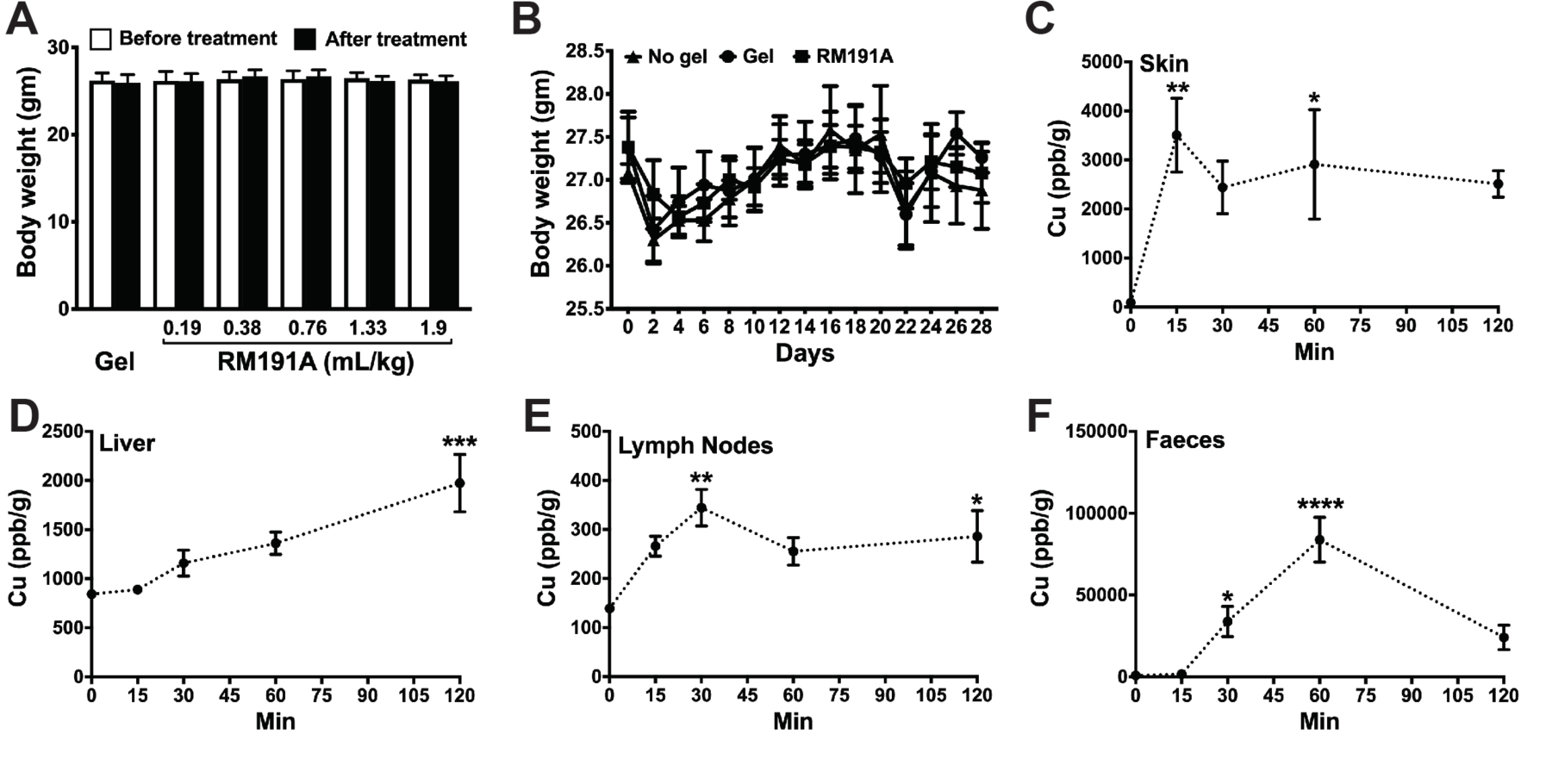
RM191A is non-toxic and readily bioavailable in mice. (A) Body weights (g) of mice treated with different topical doses of RM191A gel or gel (control) for 4 days (n = 5). (B) Body weights (g) in mice treated with topical doses of RM191A (0.19 mL/kg) or gel for 29 days (n = 4 - 9). Copper levels in (C) skin, (D) liver, (E) lymph nodes and (F) faeces over time measured using ICP-MS (n = 4 per time point). Data expressed as mean ± SEM. *p < 0.05, **p < 0.005, ***p < 0.0005 and ****p < 0.00005 by One-way ANOVA.

Next, we studied if RM191A was also well tolerated over long-term treatment. As previously, the dorsal area of 8 week old mice was shaved, and the animals were either left untreated or treated topically daily with 50 µL of gel (control) or RM191A at 0.19 mL/kg body weight for 29 days. No significant changes in body weight, water intake or food intake were observed between the three groups (Fig. 4B, S4A). The fat content, lean mass and water content, as measured using EchoMRI, were similar among these three groups (Fig. S4B). After confirming that RM191A treatment did not have any adverse impact on the general health of mice, we explored if its exposure altered the behaviour of mice. In a standard rotarod test, the animals from all three groups performed equally well, indicating no significant impact on the motor coordination in mice (Fig. S4C). Both gel and RM191A-treated groups exhibited higher spontaneous rearing behaviour compared to the untreated group (Fig. S4D). Although this increase was not significant, it implied that the gel might improve exploratory behaviour of mice. In order to confirm that there was no overt toxicity in the tissue of various organs, wet tissue weights were measured, and no differences between treated and control groups were observed (Table S5). Furthermore, histological analyses of these tissues by hematoxylin and eosin (H&E) staining showed normal morphology across all groups (Fig. S4E).

Following up on the results from the toxicity studies and skin explant experiment, we further explored RM191A’s ability to penetrate through the skin into the body. The primary goal was to determine whether or not RM191A remained on the skin where it could be washed away, or if it penetrated through the skin into the blood. To gain these insights, we applied 50 µL of RM191A (1.9 mL/kg body weight) topically to the shaved dorsal surfaces of 8 week old mice and measured total copper content in different animal tissues at different time points using ICP-MS. As expected, the amount of copper in the skin steadily increased within 15 minutes and stayed steady upto 2 hrs following RM191A treatment (Fig. 4C). There was also a significant increase in copper levels in internal tissues, including liver, lymph nodes and kideney (Fig. 4D, 4E, S4F), which implied that topically applied RM191A was able to penetrate through the skin layers and into the bloodstream (Figure S4G). The ICP-MS results demonstrated that the copper in RM191A follows the common mammalian metabolic route, with most of the copper being metabolized by the liver, the remainder by the kidneys and was excreted within one hour via the faeces (Fig. 4F) and urine (Fig. S4H). Moreover, the copper was cleared from the body within 24 hrs as its levels in tissues and blood collected 24 hrs after RM191A application reduced to the same level as the control group at this time point (Table 3).

**Table 3:**
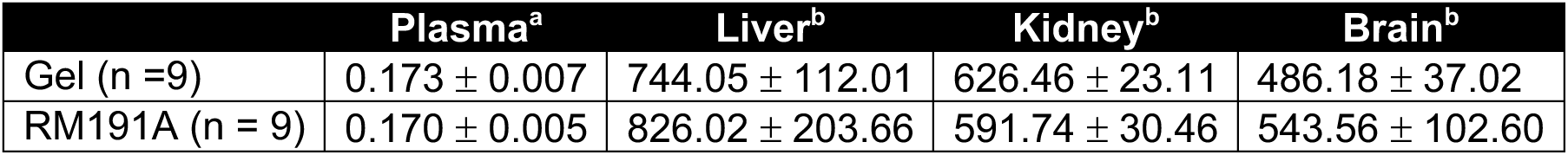
Copper contents (ppb/µL^a^ or ppb/g^b^) in plasma and tissues 24 hours after RM191A or gel treatment in mice.

The changes in brain copper levels indicated that RM191A might be able to cross the blood-brain-barrier (Fig. S4I). This was further investigated using a standard *in vitro*model for human blood-brain-barrier transmission which demonstrated that 50.79% of RM191A at 0.4 mg/mL concentration crossed the BBB in 2 hrs, and 95.65% crossed the BBB within 6 hrs (Fig. S4J).

### RM191A attenuates inflammation and improves wound healing in mice

Our previous *in vitro*data demonstrated that RM191A exhibits anti-inflammatory activity via suppression of ROS overproduction and the beneficial modulation of various signalling pathways. To test this *in vivo*, we employed a TPA (12-O-tetradecanoylphorbol-13-acetate)-induced mouse ear edema model to evaluate the anti-inflammatory activity of RM191A in mice (48). TPA treatment doubled the ear weight after 6 hrs when compared to vehicle (acetone)-treated control (Fig. 5A). Pre-treatment with RM191A (50 µL of 21 µg/mL) reduced the ear weight by 83% compared to gel and TPA-treated control. H&E staining of ear cross-sections showed increased ear thickening and leukocyte infiltration in TPA and gel-treated group, which was significantly attenuated in the RM191A pre-treated group.

**Figure 5:**
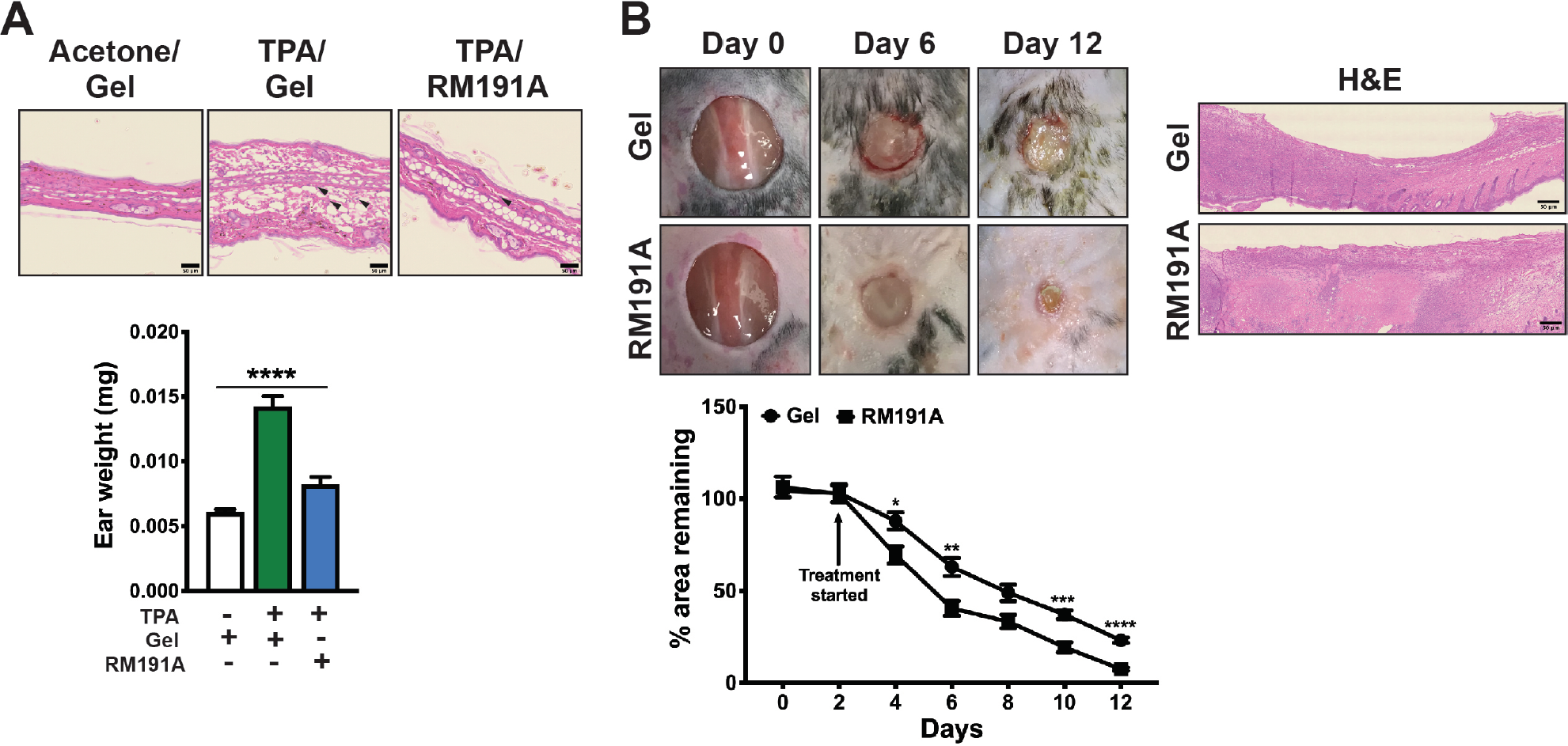
RM191A attenuates inflammation and improves wound healing in mice. (A) Representative H&E staining images of ear sections and ear weights from mice pre-treated with RM191A or gel, and then with TPA (n = 6). The arrowheads indicate the infiltrated leukocytes. (B) Representative images of wound areas and percentage wound closure in mice treated with RM191A or gel (n = 9). The representative H&E staining images of wound sections on Day 12 are shown in the right. Data expressed as mean ± SEM. *p < 0.05, **p < 0.005, ***p < 0.0005 and ****p < 0.00005 by t-test or One-way ANOVA.

Finally, we tested the effect of RM191A in wound regeneration in mice. A circular wound of 10 mm diameter was created on the dorsal surfaces of 12 week old male mice, after which 50 µL of RM191A (21 µg/mL) or gel alone was topically applied every 2 days to the wound area. Only two days after the start of RM191A treatment, 31% wound regeneration was observed in RM191A-treated group compared to only 12% wound closure in the control group (Fig. 5B). After 12 days, 93% of the skin was regenerated in the RM191A-treated animals, whereas 77% of the wound area was healed in gel-treated mice. H&E staining of the wound areas at 12 days showed regeneration of both epidermis and dermis layers in RM191A-treated group, whereas incomplete restoration of skin layers in the gel-treated group.

## Discussion

Native SODs are protective against various oxidative stress-mediated skin disorders. This fact has invoked interest in SOD mimetics for therapeutic applications. Among the various classes of SOD mimetics, only manganese porphyrin complexes have shown promise in clinical trials (49,50). This is likely due to the fact that unlike previously reported iron and copper coordination complexes, manganese-based SOD mimetics have lower toxicity, higher activity and increased stability. RM191A is the next generation copper-based SOD mimetic that is highly stable and whose activity surpasses native SOD’s abilities to scavenge free radicals, and exceeds most of the manganese-based SOD-mimetics. There are several reasons for RM191A’s high SOD-like activity. Each molecule of RM191A can react with three O_2_^-^ ions while reducing Cu^2+^-Cu^3+^ dipole to Cu^+^-Cu^+^, whereas Mn^2+^-based SOD-mimetics can react with one O_2_^-^ ion (Fig. S2E). Moreover, the reduction potential indicates that its k_cat_ for Cu^3+^/Cu^2+^ reduction is likely to be higher than that of Cu^2+^/Cu^+^ reduction, although this entails further investigation. Moreover, in cells RM191A can upregulate various antioxidant genes, such as heme oxygenase, heat shock protein and thioredoxin reductase, which can further boost its antioxidant capacity.

Despite its water solubility, RM191A was able to penetrate the skin and distribute into different internal organs. Our toxicity results demonstrated that RM191A is well-tolerated in animals. A 26 g mouse can tolerate up to 50 µL of topical RM191A with no side effects. This is equivalent to 144 mL applied to the skin of a 75 kg human. We have further found that RM191A is stable in light and heat up to 30°C for more than 12 months and that the RM191A remains active and safe in a topical hydrogel formulation over this timeframe. As far as we are aware, all previous work identifying the biological activities of copper, including its molecular mechanisms, its role in oxidative stress, and its homeostasis, specifically relate to Cu^2+^ as it would normally be present in the body (51-54). Therefore, the biological activity of a Cu^3+^ as present in RM191A is reported here for the first time.

Topical RM191A showed promise as a photoprotective agent in human skin and was at least as active as 1,25D, generally acknowledged for its photoprotective properties (34,45-48). One of the obvious reasons for this is the reduction of ROS and RNS by RM191A, although their levels were not directly tested in the explants. This may even explain the lower CPD after 3 hrs, since ROS and RNS damage DNA repair enzymes and thus inhibit DNA repair (55-57). RM191A may reduce DNA damage by other mechanisms, such as increasing energy available for DNA repair, upregulating DNA repair enzymes or enhancing mitochondrial repair (34). However, to verify this and other results it will be necessary to repeat this study in explants prepared from a range of donors with different skin types. Inadequately repaired DNA damage is a key factor leading to mutations and then to UV-induced skin tumours as well as photo-aging (45). It would be worth testing whether RM191A application in a chronic UV-exposure model reduces skin tumours and/or photo-aging in the UV exposed areas.

The topical application of the RM191A demonstrated remarkable inhibition of inflammation in the TPA mouse ear model, which has significant implications for human use for the treatment of chronic inflammatory-driven diseases including skin conditions like psoriasis (58). Similar to LPS, ROS generated by TPA can activate the inflammatory cascade (59), which we have found is significantly inhibited by RM191A. The direct involvement of RM191A in downregulating inflammatory pathways was corroborated by gene expression analysis in NHDF, where RM191A inhibited the expression of many genes reported to be involved in skin inflammation. It is possible that anti-inflammatory properties of RM191A are partly mediated by its ROS-scavaging activitiy. Although RM191A treatment had marked improvement in the healing rate of large, full-thickness excisional wounds in mice, the mechanism is not very clear to us as wound healing response is known to be orchestrated by retrograde inflammation and ROS generation (60,61). It is possible that RM191A facilitates the formation of new tissues by upregulating key repair genes and also reduces unwanted inflammation during the healing process. Nonetheless, it will be interesting to investigate how RM191A performs in chronic non-healing wounds and diabetic ulcers. RM191A’s ability to reduce oxidative stress, inflammation and modulate pain-asscoiated biomarkers indicate that RM191A may have the potential to attenuate chronic pain. Although measurement of pain was beyond the scope of this study, RM191A is currently the subject of a human clinical trial for the treatment of neuropathic pain (HREC/16/HAWKE/483).

In addition to demonstrating robust antioxidant and anti-inflammatory properties, RM191A might also facilitate the incorporation of copper into SOD1 and SOD3, ceruloplasmin and tissue repair enzymes - lysyl oxidase and elastase, which in turn can accelerate healing and inhibit inflammation. A similar mechanism has been proposed for Cu-ATSM, which is currently in clinical trial for motor neurone disease (MND) (62,63). It may also be that RM191A is protective of mitochondria by eliminating ROS before it can cause mitochondrial damage, effectively preserving normal cell signalling and maintaining the integrity of the mitochondria.

Finally, oxidative stress has been linked to a multitude of human diseases and conditions, including aging, bipolar disorder, cancer, chronic morphine intolerance, diabetes, eye diseases, fibromyalgia, ischemia, pain, radiation injury, reperfusion-related injuries – for example heart attack, stroke, and organ dysfunction; as well as neurodegenerative disorders like Alzheimer‘s, Parkinson‘s, MND, multiple sclerosis and epilepsy, to mention only a few (64). RM191A’s ability to almost instantly neutralize ROS, simultaneously diminish or eliminate the inflammatory response and easily penetrate into the brain, sets it apart as a new therapeutic and means that it likely has significant benefit in the treatment of a wide variety of human diseases/conditions.

## Conclusion

RM191A is a novel small molecule SOD mimetic that demonstrated significant SOD-like antioxidant and anti-inflammatory activities. These combined actions are likely the result of its unique, stable Cu^2+^-Cu^3+^ dipole. RM191A exhibited protective effects towards wide range of cells against oxidative stress by suppressing ROS levels and via modulating the expression of several key genes associated with oxidative stress and inflammation. The topical application of RM191A beneficially and significantly reduced three types of UV-induced DNA damage in human skin explants in the epidermis and some in dermis. This reduction in DNA damage was similar to that produced by a well-established photoprotective agent, 1,25D. RM191A easily penetrated the skin layers and was readily distributed systemically throughout the body in mice. Both short-term and long-term dosing of RM191A did not exhibit any adverse effects in mice. It’s topical application significantly reduced TPA-induced ear inflammation and accelerated excisional wound repair in mice. Therefore, RM191A is new class of SOD mimetic which exhibited beneficial effects in various skin disorders.

## Acknowledgements

This paper is dedicated to our co-author Dr. Alistair J. Laos, a brilliant young mind and a promising scientist, whom we lost too soon. The mouse experiments were supported by NSW-Government Tech Voucher Grant awarded to RR MedSciences and AD.

---

**Table S1:** Fold change in key oxidative stress gene expression in RM191A-treated NHDF compared to control.

| Gene | Fold Change |
| --- | --- |
| HMOX1 (Heme oxygenase (decycling) 1) | 67.56 |
| SLC7A11 (Solute carrier family 7 (anionic amino acid transporter light chain, xc- system), member 11) | 8.53 |
| HSPA1A (Heat shock 70kDa protein 1A) | 7.45 |
| SRXN1 (Sulfiredoxin 1) | 6.18 |
| GCLM (Glutamate-cysteine ligase, modifier subunit) | 5.75 |
| TXNRD1 (Thioredoxin reductase 1) | 5.27 |
| SQSTM1 (Sequestosome 1) | 2.19 |
| NQO1 (NAD(P)H dehydrogenase, quinone 1) | 2.02 |
| EPX (Eosinophil peroxidase) | -2.28 |
| NOX4 (NADPH oxidase 4) | -3.90 |

Fold regulation cut off = 2, p-value cut off = 0.05

**Table S2:** Fold change in key inflammatory gene expression in RM191A-treated NHDF compared to control.

| Gene | Fold Change |
| --- | --- |
| IL8 (Interleukin 8) | 6.31 |
| CCL26 (Chemokine (C-C motif) ligand 26) | 4.25 |
| TLR4 (Toll-like receptor 4) | 2.30 |
| CCL7 (Chemokine (C-C motif) ligand 7) | -2.19 |
| CXCR6 (Chemokine (C-X-C motif) receptor 6) | -2.80 |
| CCRL1 (Chemokine (C-C motif) receptor-like 1) | -3.30 |
| CMKLR1 (Chemokine-like receptor 1) | -6.18 |

Fold regulation cut off = 2, p-value cut off = 0.05

**Table S3:** Fold change in key pain gene expression in RM191A-treated NHDF compared to control.

| Gene | Fold Change |
| --- | --- |
| PLA2G1B (Phospholipase A2, group IB (pancreas)) | 2.32 |
| PTGS2 (Prostaglandin E synthase 2) | -2.10 |
| PTGER1 (Prostaglandin E receptor 1 (subtype EP1), 42kDa) | -2.16 |
| IL18 (Interleukin 18 (interferon-gamma-inducing factor)) | -2.25 |
| ADORA1 (Adenosine A1 receptor) | -2.57 |
| EDNRA (Endothelin receptor type A) | -2.64 |
| KCNIP3 (Kv channel interacting protein 3, calsenilin) | -3.04 |
| HTR2A (5-hydroxytryptamine (serotonin) receptor 2A) | -7.00 |

Fold regulation cut off = 2, p-value cut off = 0.05

**Table S4:**
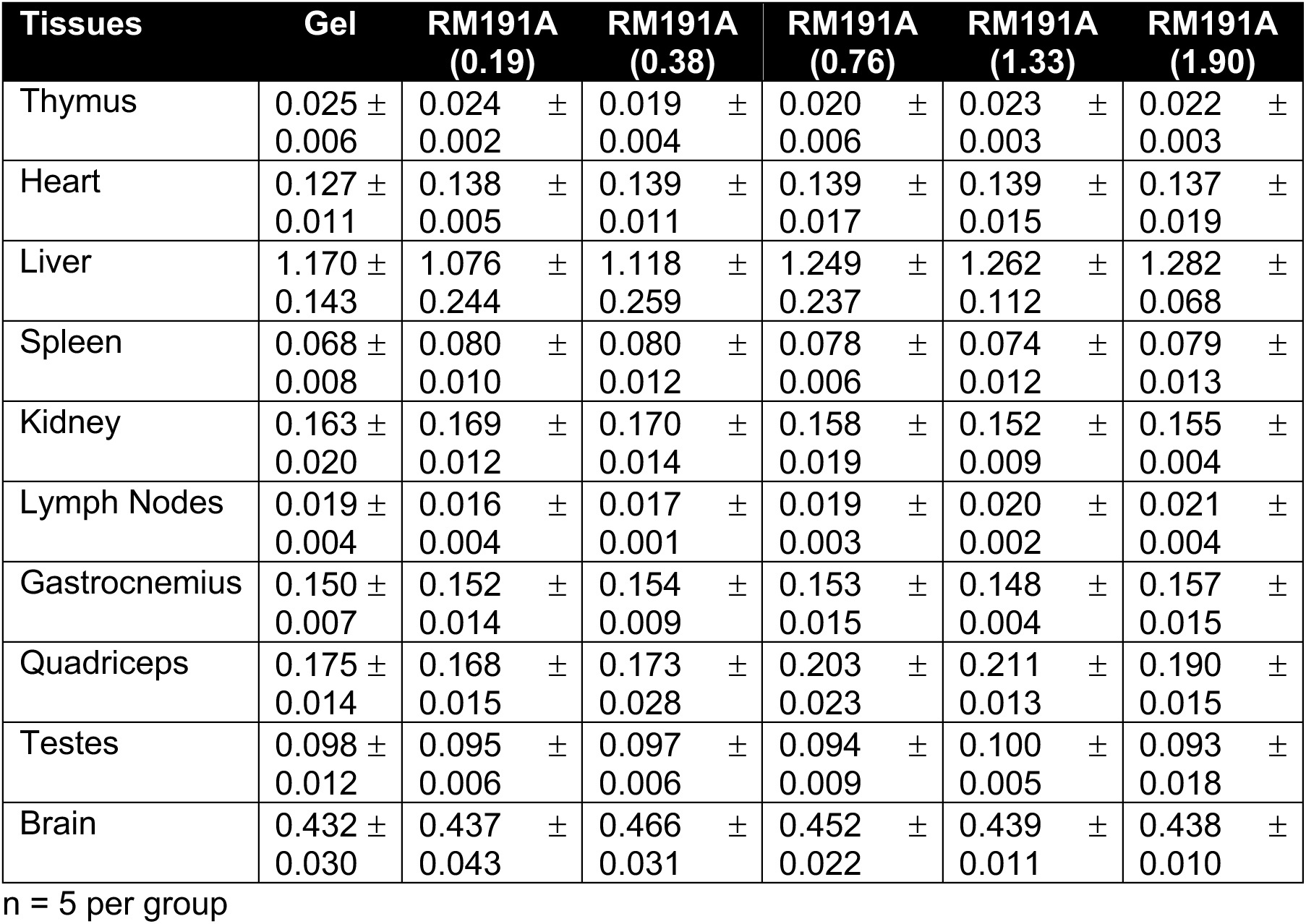
Tissue weights (g) in mice treated with different topical doses (mL/kg) of RM191A or gel (control) for 4 days.

**Table S5:** Tissue weights (g) in mice treated with topical doses of RM191A (0.19 mL/kg body weight) or gel (control) for 29 days.

| Tissues | No gel (n = 4) | Gel (n = 9) | RM191A (n = 9) |
| --- | --- | --- | --- |
| Thymus | 0.025 ± 0.003 | 0.025 ± 0.003 | 0.023 ± 0.005 |
| Heart | 0.131 ± 0.008 | 0.141 ± 0.016 | 0.139 ± 0.014 |
| Liver | 1.123 ± 0.052 | 1.213 ± 0.115 | 1.156 ± 0.136 |
| Spleen | 0.074 ± 0.003 | 0.067 ± 0.007 | 0.069 ± 0.013 |
| Kidney | 0.152 ± 0.012 | 0.167 ± 0.016 | 0.174 ± 0.011 |
| Lymph Nodes | 0.020 ± 0.004 | 0.018 ± 0.004 | 0.016 ± 0.004 |
| Gastrocnemius | 0.160 ± 0.004 | 0.162 ± 0.009 | 0.163 ± 0.012 |
| Quadriceps | 0.193 ± 0.023 | 0.195 ± 0.023 | 0.192 ± 0.022 |
| Testes | 0.113 ± 0.007 | 0.106 ± 0.009 | 0.104 ± 0.006 |
| Brain | 0.413 ± 0.025 | 0.435 ± 0.015 | 0.448 ± 0.024 |

**Figure S1:**
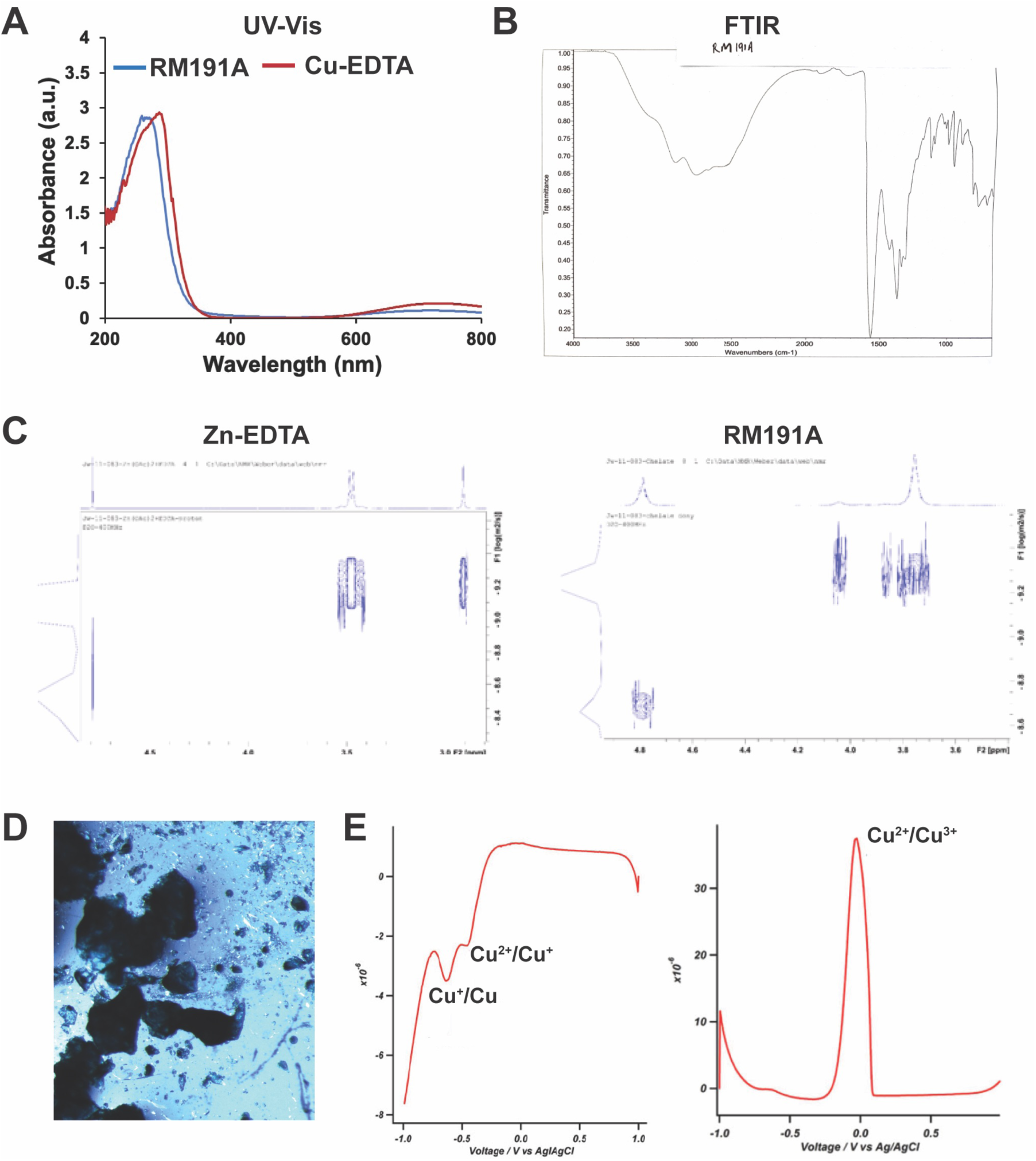
Synthesis and characterization of RM191A. (A) UV-visible spectrum of RM191 and Cu-EDTA complex. (B) FTIR spectrum of RM191A. (C) DOSY 2D spectrum of EDTA-Zinc complex and RM191A in D_2_O. (D) White light micrograph of RM191A crystals. (E) Cyclic Voltammetry results of RM191A showing two reduction peaks and one oxidation peak.

**Figure S2:**
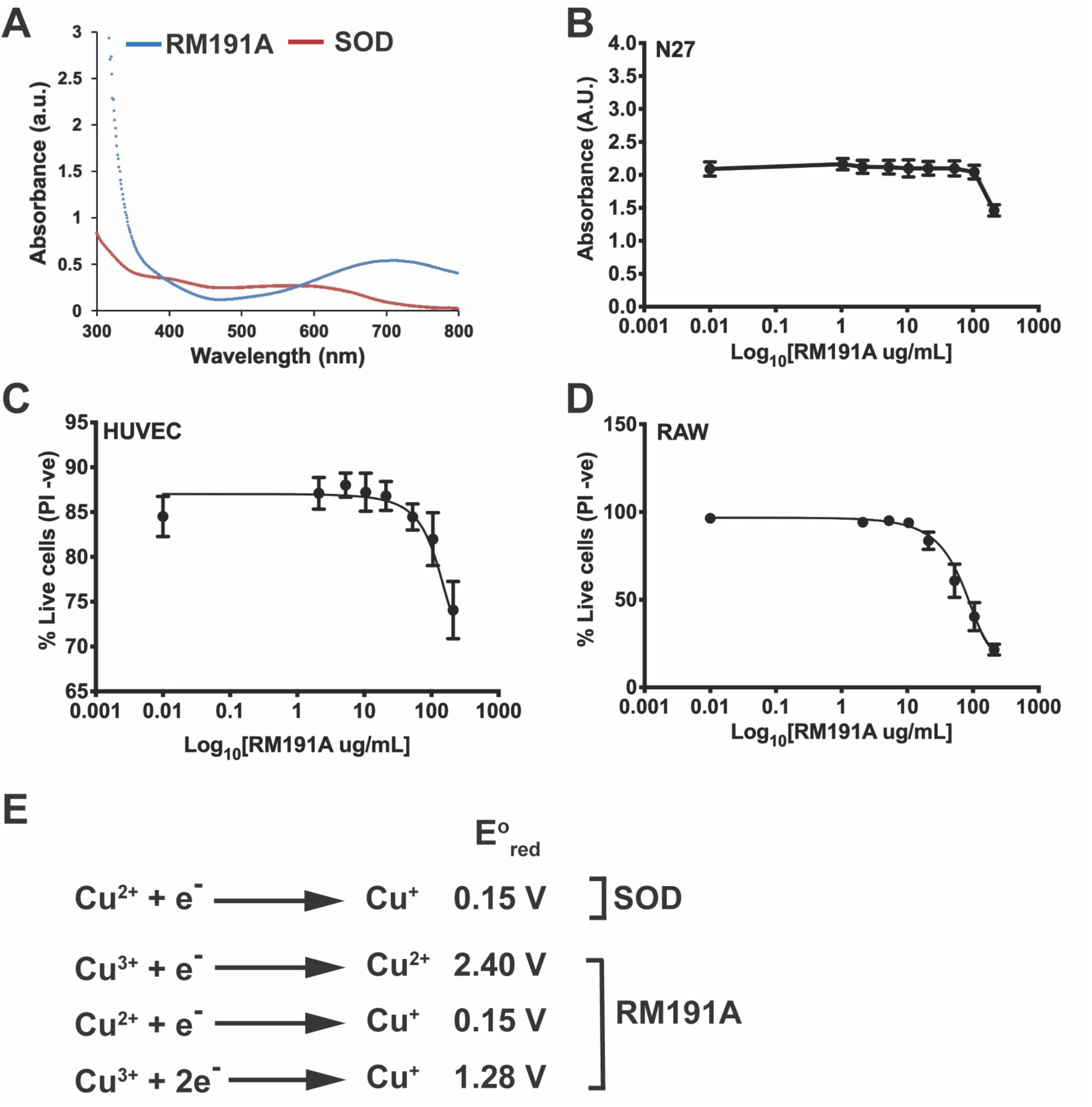
RM191A has potent antioxidant and anti-inflammatory properties. (A) UV-visible spectrum of RM191 and bovine SOD1. (B) Dose dependent curve of RM191A for N27 cells, as measured by MTT assay (n = 8). (C) Dose dependent curve of RM191A for HUVEC, as measured by PI exclusion (PI -ve) in flow cytometry (n = 6). (D) Dose dependent curve of RM191A for RAW 264.7 cells, as measured by PI exclusion (PI -ve) in flow cytometry (n = 6). (E) Reduction potential of Cu at various oxidative states present in SOD and RM191A.

**Figure S3:**
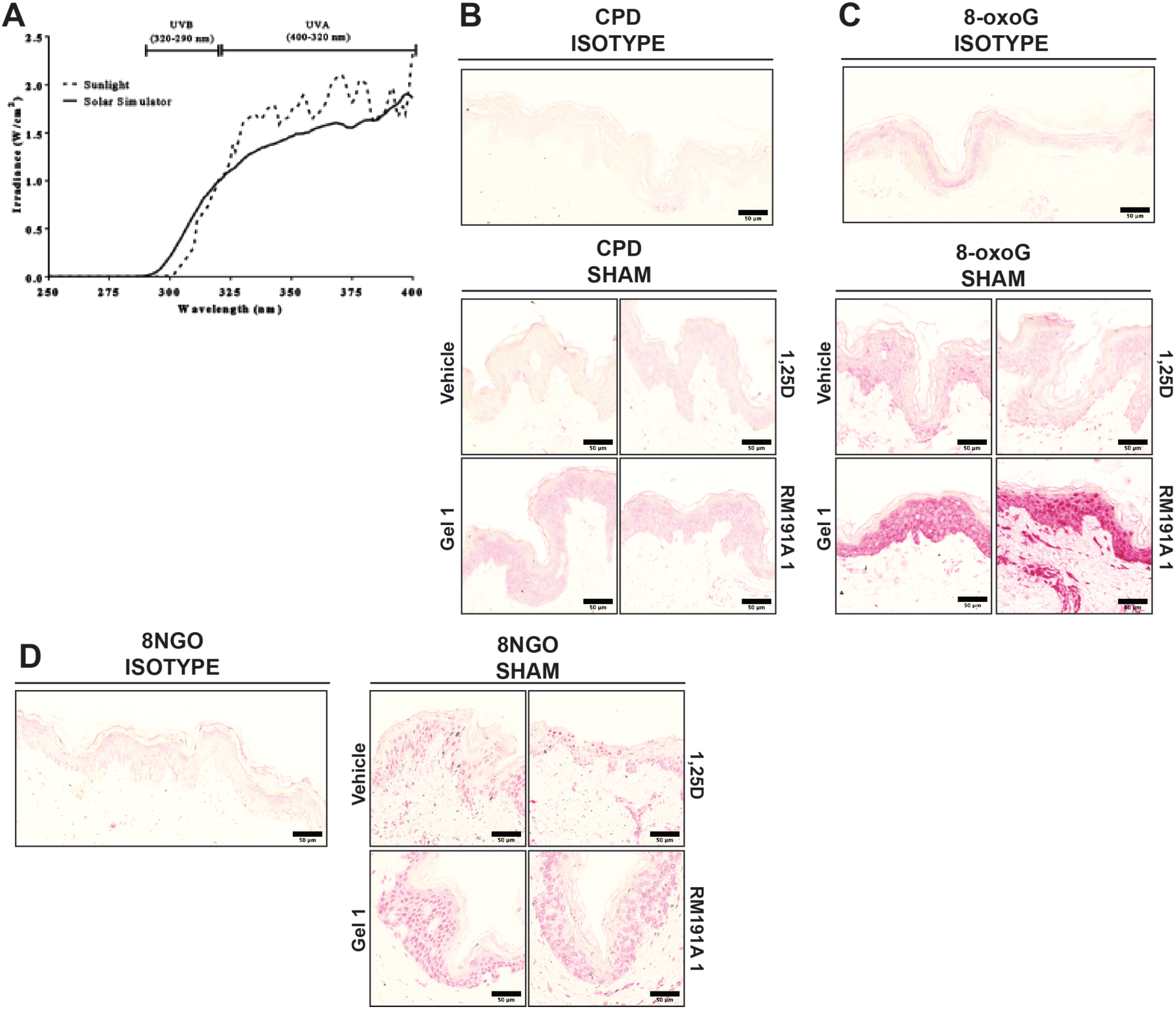
RM191A protects against UV-induced oxidative stress. (A) A comparison of the output of natural sunlight with the Oriel 450W solar simulator containing an atmospheric attenuation filter. The dotted line represents the output of natural sunlight at midday in October in Sydney, Australia, while the solid line represents the output of the solar simulator used in all experiments in human explant skin. While the output of the solar simulator closely resembles that of natural sunlight, the former emits 10% UVB, while the latter emits 5% UVB. (B) Representative images of immunohistochemical staining of CPDs in human explants from isotype control and sham exposed explants. There was little staining in the isotype control. Sham irradiated skin showed low levels of nuclear staining. (C) Representative images of immunohistochemical staining of 8-oxoG in human explants from isotype control and sham exposed explants. (D) Representative images of immunohistochemical staining of 8NGO in human explants from isotype control and sham exposed explants.

**Figure S4:**
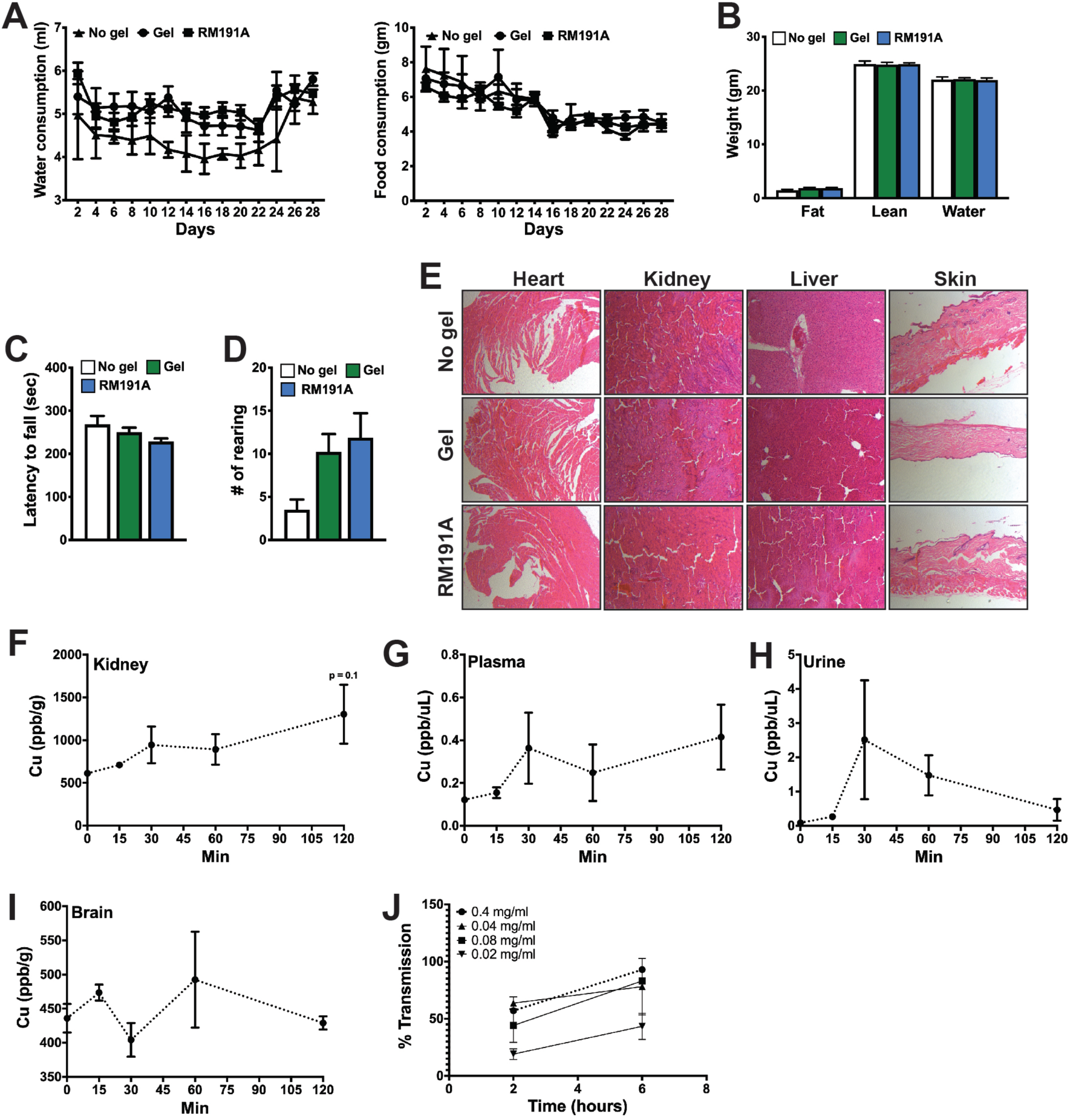
RM191A is non-toxic and readily bioavailable in mice. (A) Water and food consumption, (B) EchoMRI, (C) rotarod performance and (D) rearing behaviour of mice untreated and treated with topical doses of RM191A (0.19 mL/kg) or gel (control) for 29 days (n = 9). (E) H&E staining of heart, kidney, liver and skin from mice untreated and treated with topical doses of RM191A or gel for 29 days. Copper levels in (F) kidney, (G) plasma, (H) urine and (I) brain over time measured using ICP-MS (n = 4). (J) Percentage transmission of RM191A over time in the *in vitro* blood-brain-barrier model (n = 2).

## Graphical Abstract

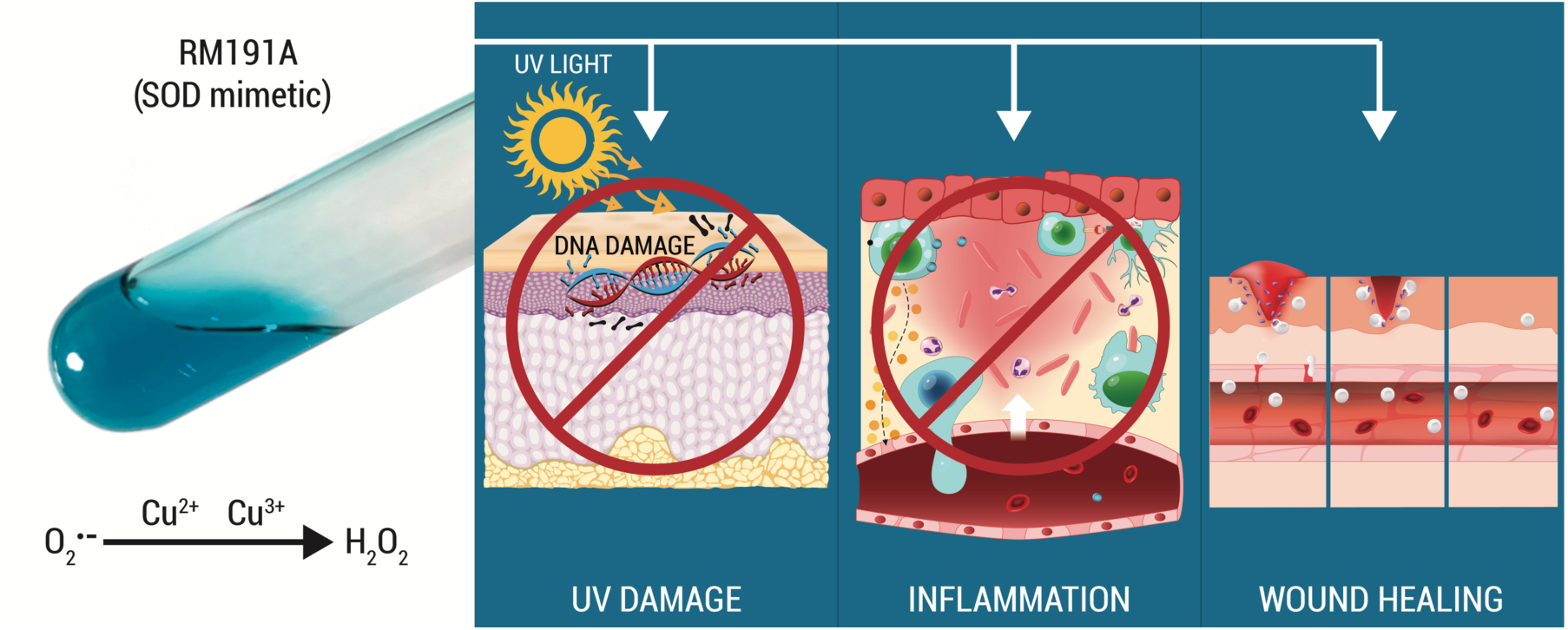

